# Hierarchical Transcriptomic and Epigenetic Recovery and Remodeling in the Developing Hippocampus Following Early-life Environmental Insults: An Iron Deficiency Rat Model

**DOI:** 10.64898/2026.07.22.740094

**Authors:** Shirelle X. Liu, Zia L. Maxim, Carrie Walls, Cleo Kilpatrick, Christopher Faulk, Michael K. Georgieff, Phu V. Tran

## Abstract

**Backgrounds:** Early-life environmental insults cause persistent neurodevelopmental abnormalities accompanied by transcriptional and epigenetic dysregulation despite removal of the original insult or postnatal intervention. However, transcriptomic and epigenomic responses to developmental insults and subsequent treatment during active neurodevelopment remain insufficiently characterized. Developmental iron deficiency (ID) provides a unique model for investigating this question because iron is an essential cofactor for TET DNA dioxygenases and developmental ID causes persistent behavioral and molecular alterations despite iron repletion.

**Results:** We integrated the hippocampal transcriptome, DNA methylome (5mC), and hydroxymethylome (5hmC) in male rats at postnatal day 15 following developmental ID and postnatal iron treatment, using Oxford Nanopore sequencing for native DNA modification profiling. Developmental ID induced substantial transcriptional and epigenetic alterations associated with synaptic function, neurodevelopment, and neuroinflammation. Postnatal iron treatment induced a hierarchical response across molecular layers: while transcriptomic alterations largely normalized, 5mC showed only partial recovery, and 5hmC showed extensive *de novo* modifications. Recovered, persistent, and newly emerged epigenetic marks were associated with increasingly specialized biological functions, from broad neurodevelopmental processes to specific pathways. Furthermore, while 5mC enrichment was associated with transcriptionally suppressed pathways, 5hmC enrichment showed weaker coupling with concurrent transcriptomic activity, suggesting epigenetic poising rather than immediate transcriptional output. MergeOmics integration identified key driver genes showing post-treatment epigenetic regulation despite transcriptional recovery.

**Conclusions:** Molecular recovery following developmental ID extends beyond transcriptomic normalization, involving persistent and extensive epigenetic remodeling. This study provides a framework for understanding molecular responses following early-life environmental insults and highlights the importance of delineating persistent regulatory reprogramming.

## Background

Early-life environmental insults, such as nutritional deficiency (1–3), chemical exposure (4–6), and inflammation (7–9), can produce persistent neurodevelopmental and behavioral abnormalities despite removal of the original insult or postnatal intervention. One plausible mechanism is altered epigenetic regulation, through which transient developmental insults produce long-lasting molecular changes (10–12). In particular, DNA methylation (5mC) and hydroxymethylation (5hmC) are dynamic epigenetic marks during brain development (13, 14) that regulate neural differentiation (15, 16), synaptic plasticity (17, 18), and long-term gene expression (19). In addition, 5mC and 5hmC have been used as biomarkers for neurodevelopmental disorders, highlighting their relative stability and potential use in early disease detection (20, 21). However, how the transcriptome, 5mC, and 5hmC respond to environmental insults and subsequently recover or remodel following intervention remain insufficiently characterized.

Developmental iron deficiency (ID) provides a unique model for investigating this question. ID is the most prevalent micronutrient deficiency worldwide, particularly during pregnancy and early childhood (22). Developmental ID has been associated with deficits in memory, motor function, social behavior, and mental health that persist despite restoration of postnatal iron status (1). Importantly, iron is an essential cofactor for many epigenetic modifiers, including the Ten-Eleven Translocation (TET) DNA dioxygenases and the Jumonji and AT-Rich Interaction Domain-containing (JARID) histone demethylases. TET enzymes convert 5mC to 5hmC and subsequent derivatives (5fC and 5caC). Thus, ID can directly affect DNA methylation (23). In our previous preclinical studies, we found that developmental ID caused persistent impairments in recognition memory in adult rats despite restoration of iron status (24). These rats also showed persistent dysregulation in transcription, chromatin accessibility, and histone modification of genes involved in synaptic plasticity, neuroinflammation, and reward circuitry (24–27). Therefore, this model is suited for investigating the dynamics of the transcriptome, 5mC, and 5hmC during developmental ID and iron restoration.

Whole-genome bisulfite sequencing is considered the gold standard for genome-wide DNA methylation profiling (28, 29). However, this method does not distinguish between 5mC and 5hmC (30). Separate quantification of 5mC and 5hmC typically requires a second approach, such as oxidative bisulfite sequencing, which involves additional chemical conversion, independent sequencing libraries, and downstream analysis pipelines (30). The harsh chemical conversions cause DNA degradation, and the additional procedures increase experimental complexity and technical variability (31, 32). In contrast, third-generation long-read sequencing technologies, such as Oxford Nanopore sequencing, directly sequence native DNA molecules and simultaneously detects both 5mC and 5hmC without chemical conversion (33). This enables integrated whole-genome analysis of both 5mC and 5hmC within a single experimental and computational workflow, improving experimental consistency.

In this study, we investigated dynamic changes in the transcriptome, 5mC, and 5hmC in the developing hippocampus of male rat offspring in response to developmental ID and postnatal iron treatment using Oxford Nanopore sequencing. Iron-sufficient (IS), iron-deficient anemic (IDA), and postnatally iron-treated IDA (TIDA) offspring were generated through dietary manipulation of the dams (Fig. 1). The TIDA group received iron treatment beginning at postnatal day (P) 7, a time point at which rodent neurodevelopment approximates that of a human full-term infant (34). The Hippocampus was analyzed at P15, during late-stage neurogenesis (35) and early neuronal differentiation (36). We characterized recovered, persistent, and newly emerged transcriptional and epigenetic alterations associated with developmental ID and postnatal iron treatment. Integrative network analysis further identified functional pathways and key driver genes associated with post-treatment epigenetic remodeling despite transcriptional recovery. Our data suggest that the transcriptome, 5mC, 5hmC recover or remodel in a hierarchical manner, and that persistent, extensive epigenetic alterations may contribute to long-term neurodevelopmental and behavioral abnormalities. Our findings provide a framework of how molecular recovery may occur following early-life environmental insults in the developing brain and highlight the importance of examining persistent regulatory changes beyond restoration of gene expression.

**Figure 1.**
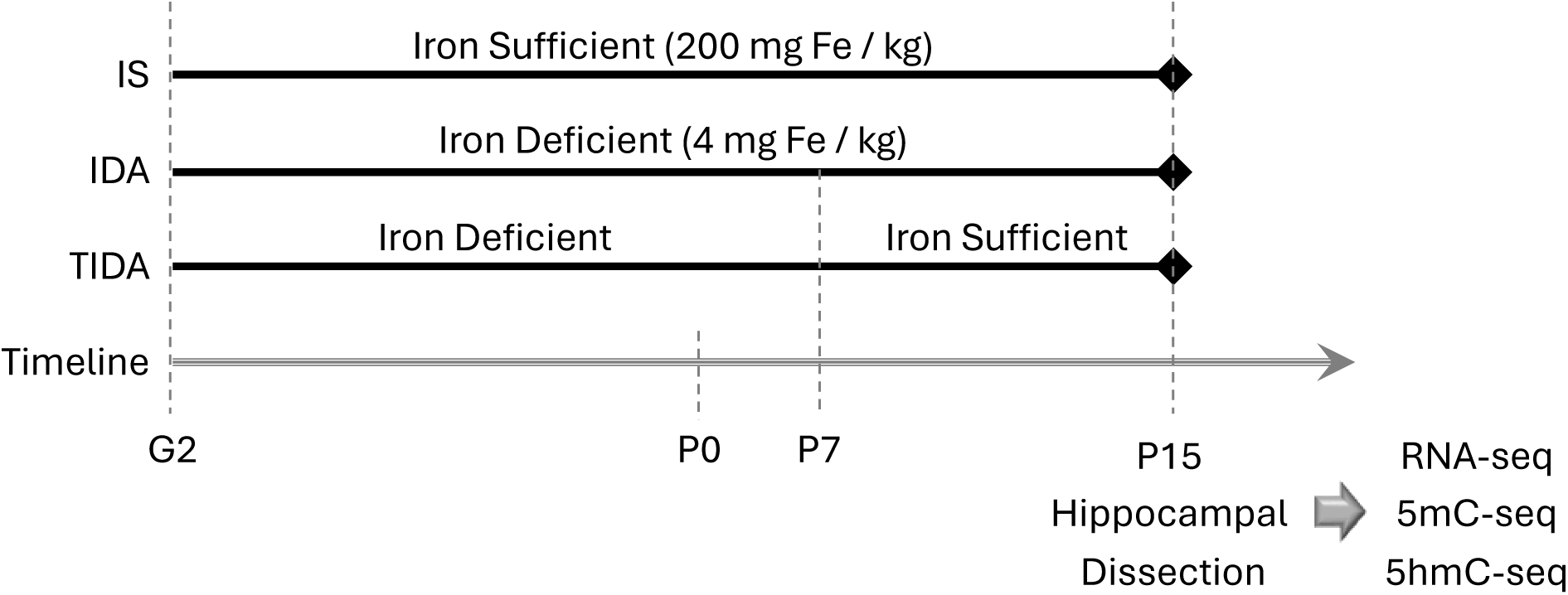
Experimental design and dietary treatments for the IS, IDA, and TIDA groups. Hippocampi were collected at P15 for RNA-seq, 5mC-seq, and 5hmC-seq.

## Results

### Transcriptome, 5mC, and 5hmC showed distinct, hierarchical patterns of recovery and remodeling following developmental ID and postnatal iron treatment

Postnatal iron treatment largely restored developmental ID-induced transcriptional alterations in the P15 male rat hippocampus. RNA-seq analysis identified 1,954 differentially expressed genes (DEGs) in the IDA vs. IS comparison (Fig. 2A). In contrast, substantially fewer (176) DEGs were identified in the TIDA vs. IS comparison, whereas 1,939 DEGs were identified in the TIDA vs. IDA comparison (Fig. 2A). An alluvial plot showed that 96.2% of the DEGs in the IDA vs. IS comparison were recovered in the TIDA vs. IS comparison (Down or Up → NS (not significant), Fig. 2B). Among the recovered DEGs, 71.6% specifically reflected the effects of postnatal iron treatment (Down → Up → NS or Up → Down → NS, Fig. 2B).

**Figure 2.**
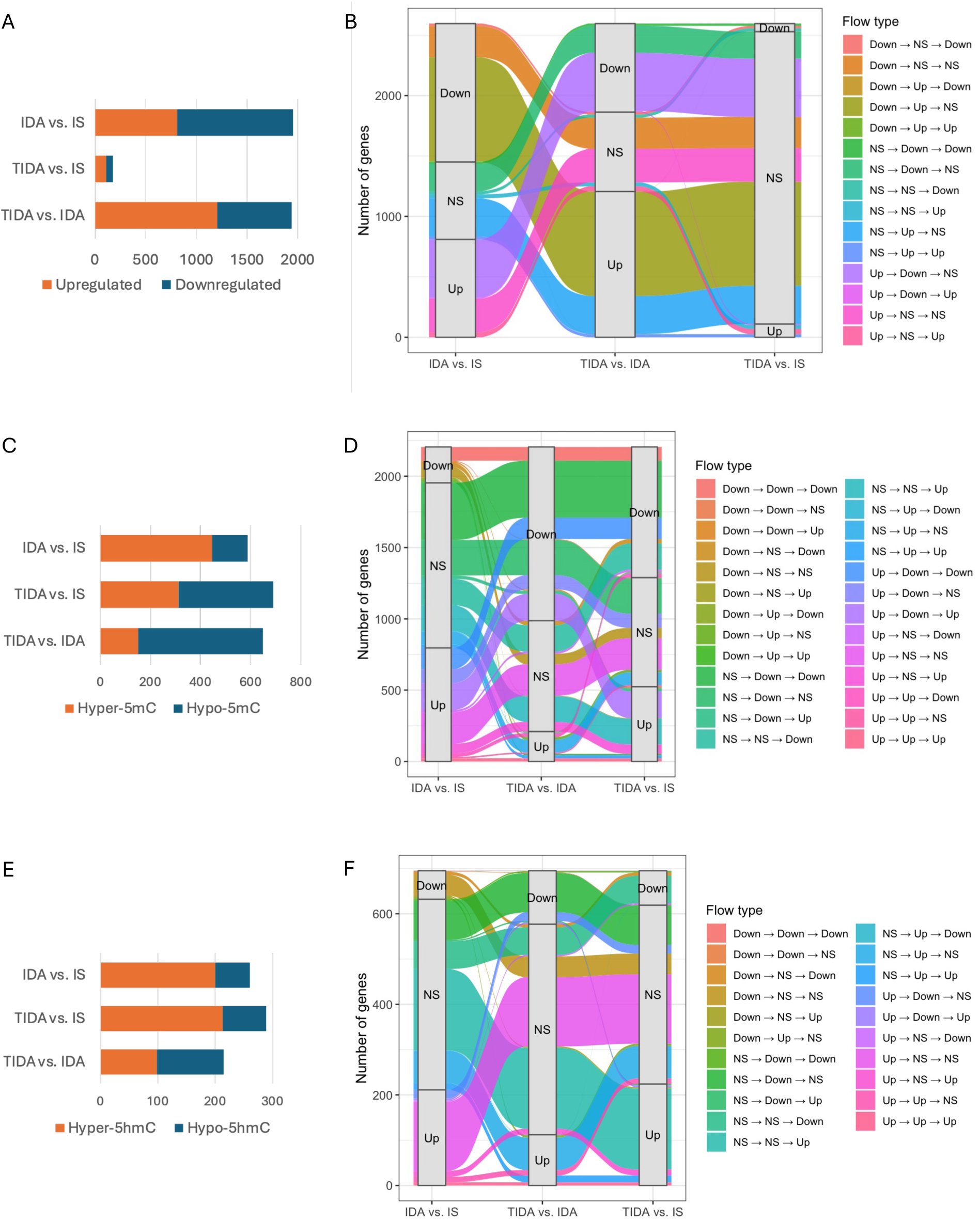
Transcriptome, 5mC, and 5hmC show distinct, hierarchical patterns of recovery and remodeling following developmental ID and postnatal iron treatment in the P15 male rat hippocampus. (A) Number of DEGs identified by RNA-seq across the IDA vs. IS, TIDA vs. IS, and TIDA vs. IDA comparisons. (B) Alluvial plot showing transitions in gene expression status across comparisons. (C) Number of DMRs identified by 5mC-seq across comparisons. (D) Alluvial plot showing transitions in 5mC enrichment at genes across comparisons. (E) Number of DhMRs identified by 5hmC-seq across comparisons. (F) Alluvial plot showing transitions in 5hmC enrichment at genes across comparisons. NS, not significant.

Postnatal iron treatment only partially restored developmental ID-induced 5mC alterations, while additional 5mC changes emerged during recovery. 5mC-seq analysis identified 588 differentially methylated regions (DMRs) in the IDA vs. IS comparison (Fig. 2C). In contrast to the transcriptome, postnatal iron treatment did not reduce the number (689) of DMRs in the TIDA vs. IS comparison, despite that 649 DMRs were identified in the TIDA vs. IDA comparison (Fig. 2C). However, an alluvial plot showed that 40.6% of the DMR-associated genes in the IDA vs. IS comparison were recovered in the TIDA vs. IS comparison (Down or Up → NS, Fig. 2D). In contrast to the transcriptome, among the recovered DMR-associated genes, only 27.5% specifically reflected the effects of postnatal iron treatment (Down → Up → NS or Up → Down → NS, Fig. 2D). In addition, 56.8% of the DMR-associated genes in the TIDA vs. IS comparison were newly emerged that were non-significant in the IDA vs. IS comparison (NS → Down or Up), among which 51.8% specifically reflected the effects of postnatal iron treatment (NS → Down → Down or NS → Up → Up) (Fig. 2D).

While postnatal iron treatment largely restored developmental ID-induced 5hmC alterations, substantial additional 5hmC changes emerged during recovery. 5hmC-seq analysis identified 261 differentially hydroxymethylated regions (DhMRs) in the IDA vs. IS comparison (Fig. 2E). Similar to the 5mC, postnatal iron treatment did not reduce the number (289) of DhMRs in the TIDA vs. IS comparison, despite that 215 DhMRs were identified in the TIDA vs. IDA comparison (Fig. 2E). However, in contrast to the 5mC, an alluvial plot showed that 86.1% of the DhMR-assocaited genes in the IDA vs. IS comparison were recovered in the TIDA vs. IS comparison (Down or Up → NS, Fig. 2F). However, among the recovered DhMR-assocaited genes, only 10.2% specifically reflected the effects of postnatal iron treatment (Down → Up → NS or Up → Down → NS, Fig. 2F). Notably, 87.3% of the DhMR-assocaited genes in the TIDA vs. IS comparison were newly emerged that were non-significant in the IDA vs. IS comparison (NS → Down or Up), among which only 6.9% specifically reflected the effects of postnatal iron treatment (NS → Down → Down or NS → Up → Up) (Fig. 2F). These results indicated that 5hmC exhibited a more complex response to iron treatment than transcriptome and 5mC.

### Overlapping DEGs across comparisons showed significant but incomplete recovery following developmental ID and postnatal iron treatment

To identify genes that were dysregulated by developmental ID, responsive to postnatal iron treatment, yet incompletely recovered, we overlapped DEGs across the 3 comparisons (Fig. 3A). Notably, all 12 overlapping genes, including *Tfrc*, a marker of functional ID, showed significant yet partial recovery (Fig. 3B). The majority of these genes are involved in hypoxia responses (*Adm*, *Ankrd37*, *Tmem45b*) or neuronal functions (*Syt8*, *Atp2a3*, *Prkcd*). Expression changes in selected genes were validated by qPCR (Fig. 3C).

**Figure 3.**
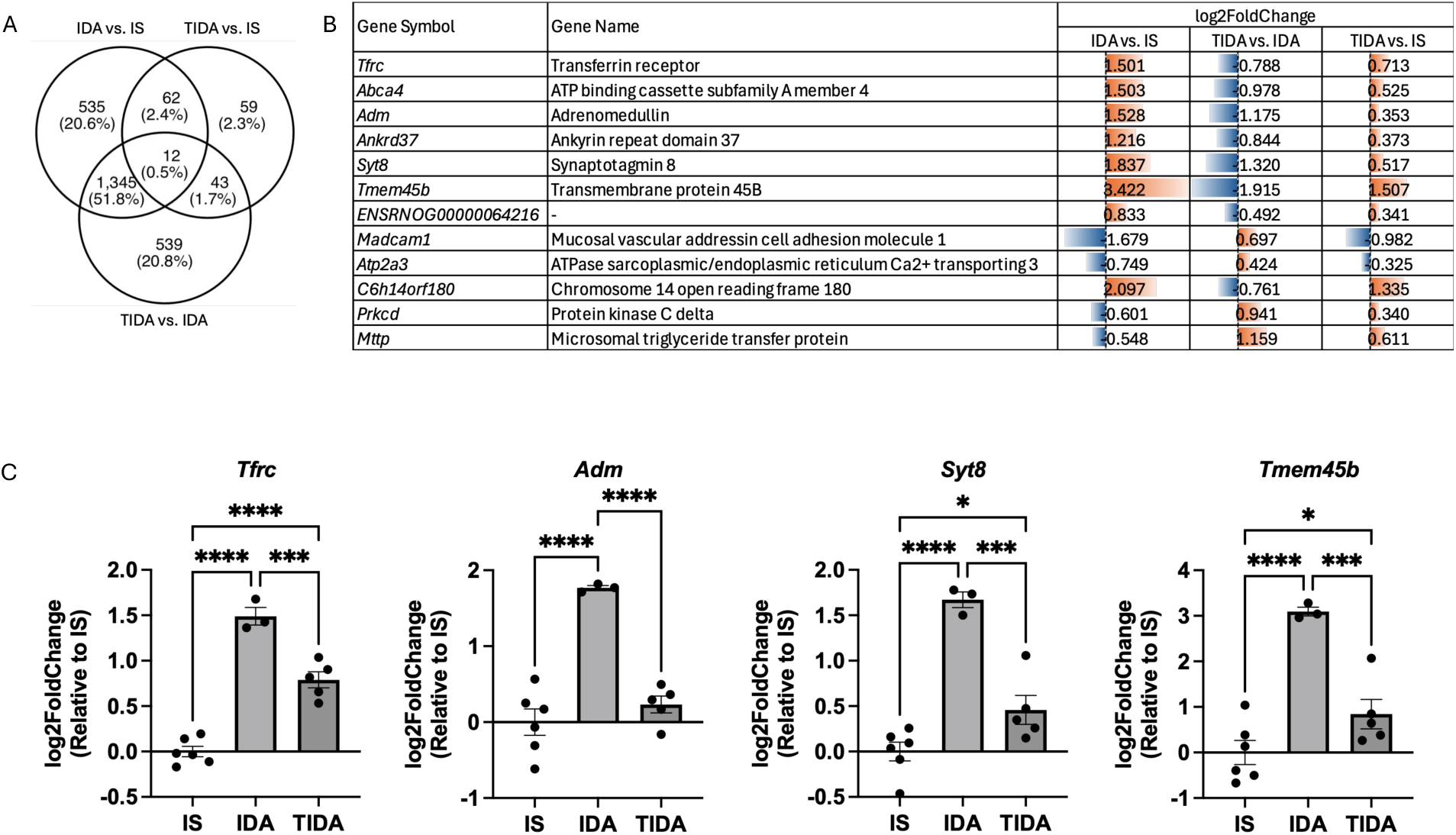
Overlapping DEGs show persistent dysregulation following developmental iron deficiency and iron treatment. (A) Venn diagram showing the overlap of DEGs across comparisons. (B) Table summarizing the 12 overlapping DEGs across the 3 comparisons that show significant but incomplete recovery. Bar lengths represent log2FoldChange; orange, upregulation; blue, downregulation. (C) qPCR validation of *Tfrc*, *Adm*, *Syt8*, and *Tmem45b*. Data are presented as mean ± SEM. n = 3–6 per group. *p < 0.05, ***p < 0.001, ****p < 0.0001.

### Recovered, persistent, and newly emerged DMRs and DhMRs were associated with increasingly specialized biological functions

A Venn diagram identified 370 recovered DMR-associated genes (exclusively in IDA vs. IS, Fig. 4A). Gene ontology (GO) analysis showed that these genes were enriched in cell migration and cell communication (Fig. 4B).144 genes associated with developmental ID-induced DMRs were non-recovered (in both IDA vs. IS and TIDA vs. IS, Fig. 4A). GO analysis showed that these genes were enriched in metabolic regulation, gene transcription, and general developmental processes (Fig. 4C). Kyoto Encyclopedia of Genes and Genomes (KEGG) analysis showed that these genes were enriched in Wnt signaling pathway (Fig. 4D). 441 genes were associated with newly emerged DMRs (exclusively in TIDA vs. IS, Fig. 4A). GO analysis showed that these genes were enriched in neurodevelopment (Fig. 4E). KEGG analysis showed that these genes were enriched in axon guidance (Fig. 4F).

**Figure 4.**
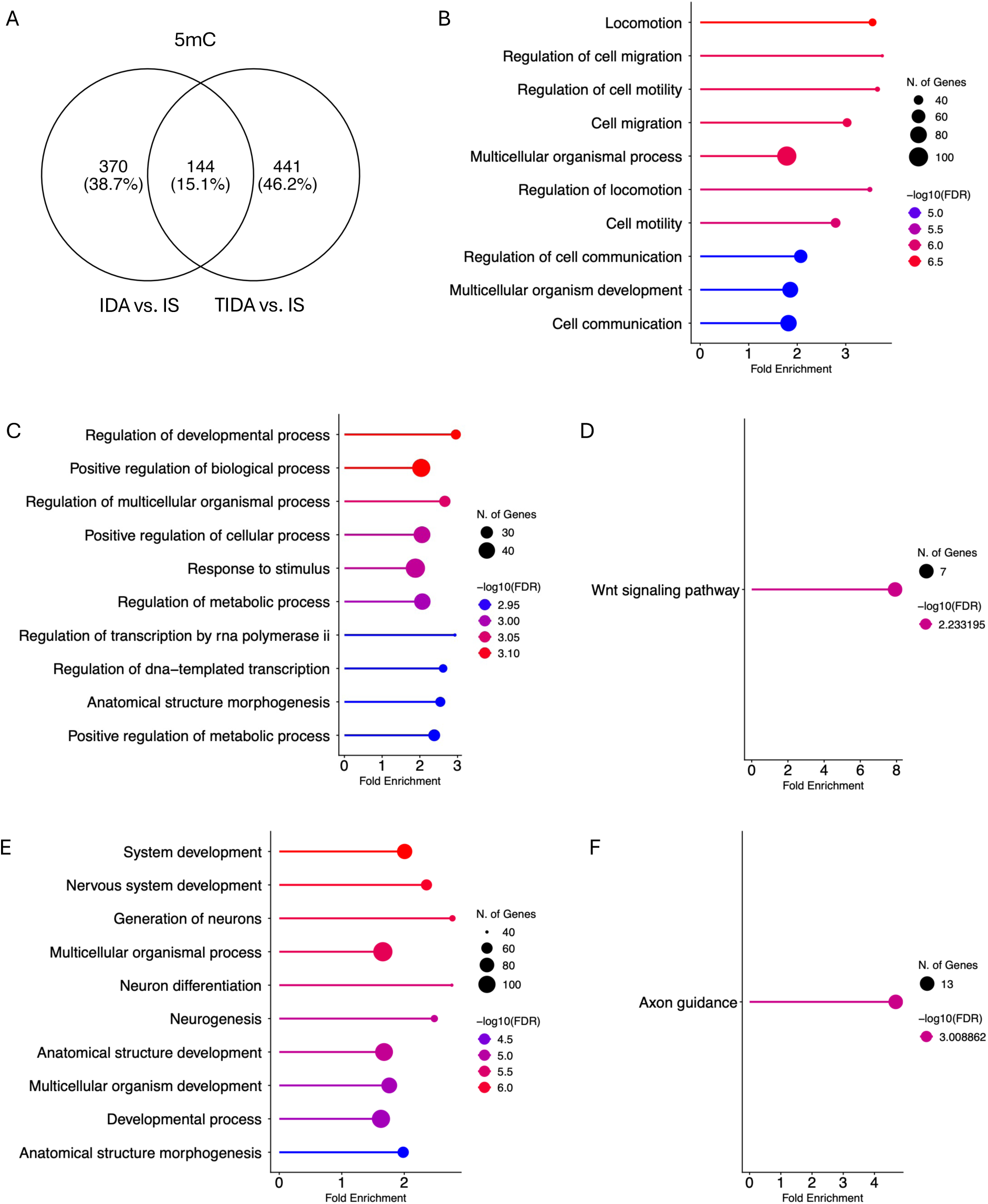
Recovered, persistent, and newly emerged DMRs are associated with increasingly specialized biological functions. (A) Venn diagram showing the overlap of DMR-associated genes between the IDA vs. IS and TIDA vs. IS comparisons. (B) GO analysis of 370 recovered DMR-associated genes (exclusively in IDA vs. IS). (C, D) GO (C) and KEGG (D) analysis of 144 non-recovered DMR-associated genes (in both IDA vs. IS and TIDA vs. IS). (E, F) GO (E) and KEGG (F) analysis of 441 newly emerged DMR-associated genes (exclusively in TIDA vs. IS).

A Venn diagram identified 223 recovered DhMR-associated genes (exclusively in IDA vs. IS, Fig. 5A). GO analysis showed that these genes were enriched in developmental and biosynthetic processes (Fig. 5B). 26 genes associated with developmental ID-induced DhMRs were non-recovered (in both IDA vs. IS and TIDA vs. IS, Fig. 5A). GO analysis showed that these genes were associated with calcium transportation (Fig. 5C). 250 genes were associated with newly emerged DhMRs (exclusively in TIDA vs. IS, Fig. 5A). GO analysis showed that these genes were associated with cell signaling and developmental processes (Fig. 5D). KEGG analysis showed that these genes were enriched in dopaminergic function and circadian entrainment (Fig. 5E).

**Figure 5.**
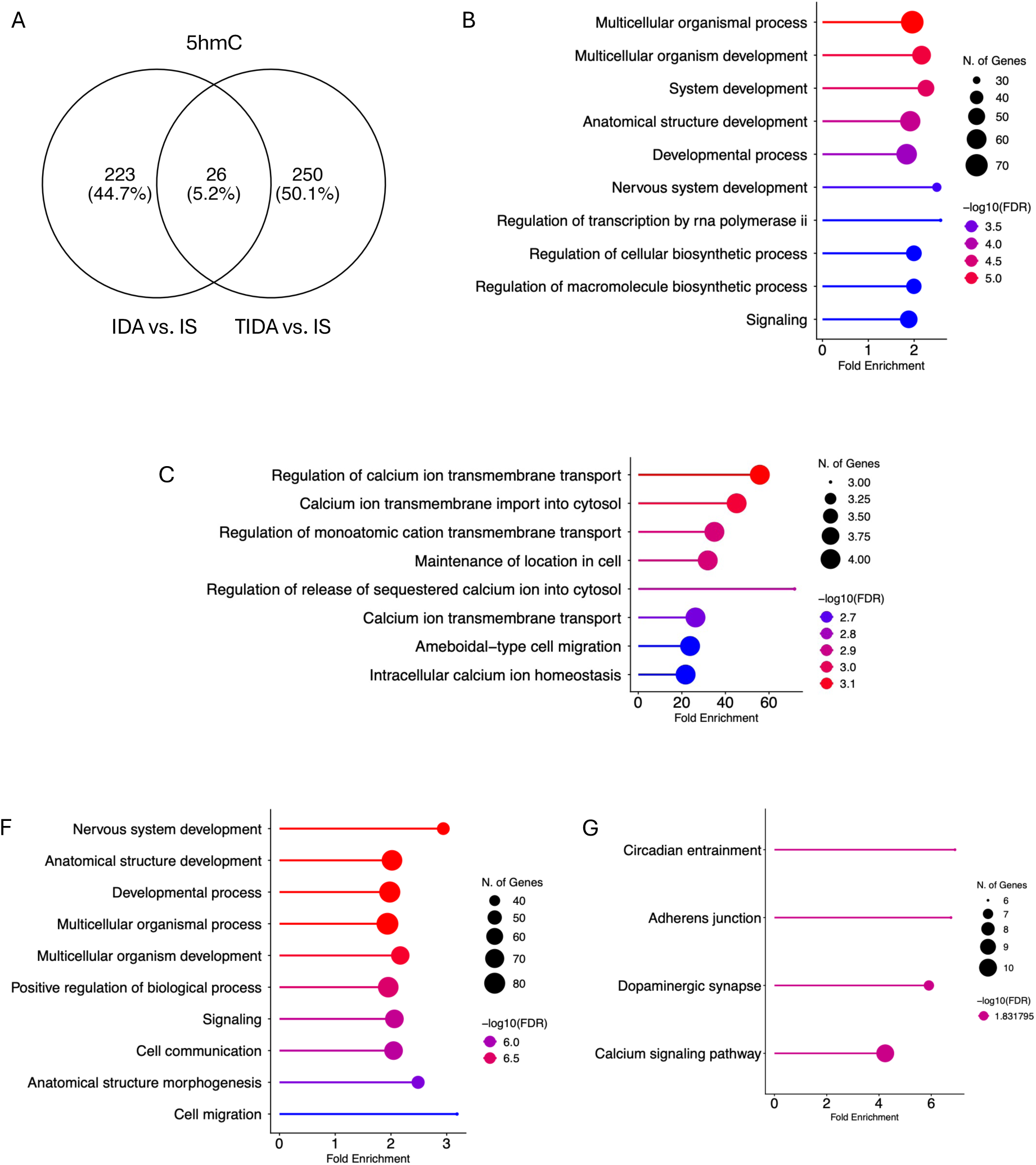
Recovered, persistent, and newly emerged DhMRs are associated with increasingly specialized biological functions. (A) Venn diagram showing the overlap of DhMR-associated genes between the IDA vs. IS and TIDA vs. IS comparisons. (B) GO analysis of 223 recovered DhMR-associated genes (exclusively in IDA vs. IS). (C, D) GO (C) and KEGG (D) analysis of 26 non-recovered DhMR-associated genes (in both IDA vs. IS and TIDA vs. IS). (E, F) GO (E) and KEGG (F) analysis of 250 newly emerged DhMR-associated genes (exclusively in TIDA vs. IS).

### Genes associated with intronic DMRs and DhMRs showed stronger enrichment for biological pathways

Since 5mC and 5hmC can exert distinct regulatory effects depending on their genomic locations, we assessed the genomic distribution of DMRs and DhMRs and their enrichment in Ingenuity Pathway Analysis (IPA) canonical pathways. DMRs and DhMRs were predominantly located within intronic and intergenic regions across the 3 comparisons (Fig. 6A, B). However, only genes associated with intronic DMRs and DhMRs showed strong enrichment in IPA canonical pathways (Fig. 6C–F). These findings suggest that intronic 5mC and 5hmC alterations may have greater functional relevance than intergenic changes in developmental ID and the response to postnatal iron treatment.

**Figure 6.**
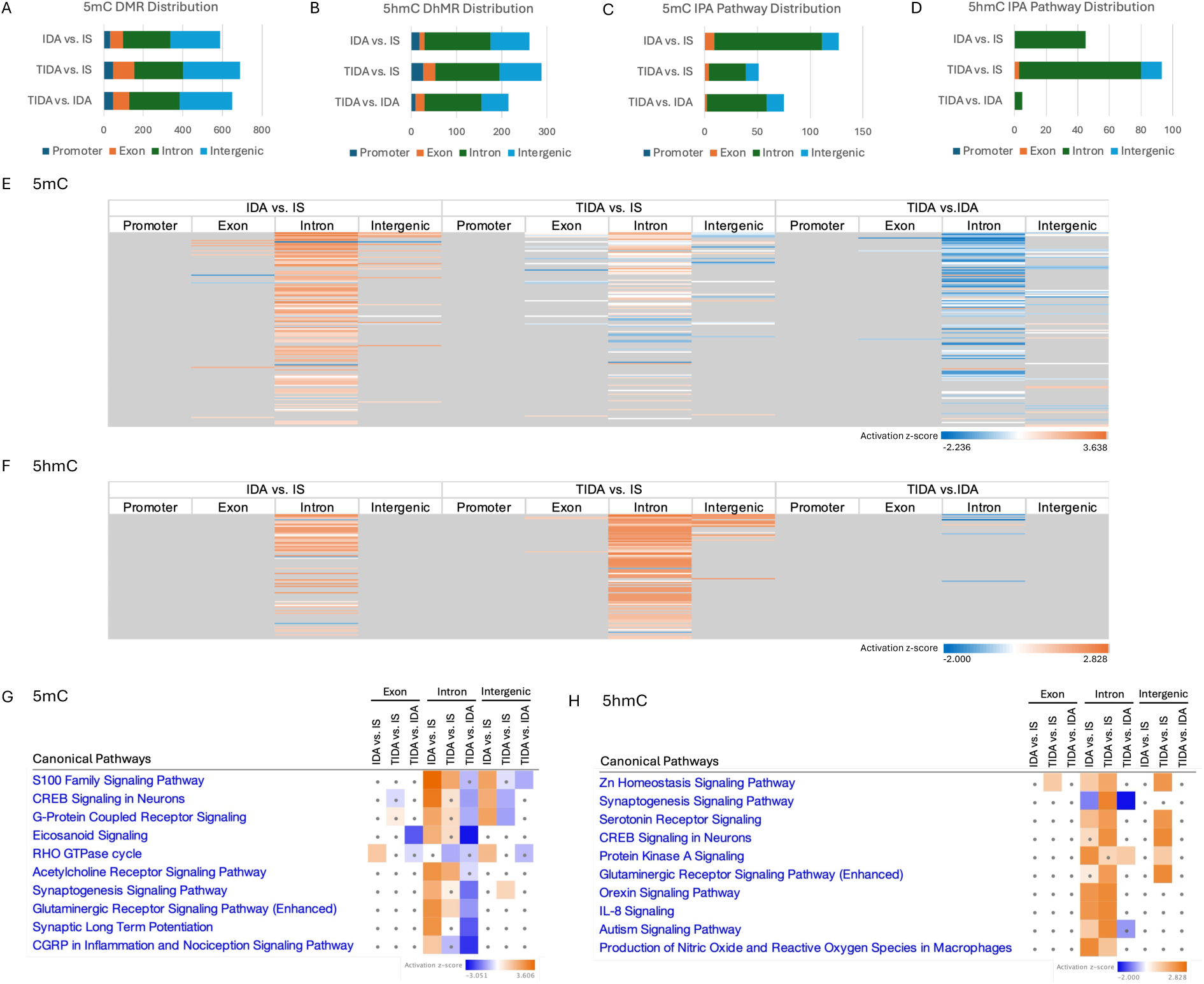
Regional distribution of DMRs and DhMRs across the genome and enriched IPA canonical pathways. (A, B) Numbers of DMRs (A) and DhMRs (B) located within promoter, exon, intron, and intergenic regions across comparisons. (C, D) Numbers of IPA biological pathways that showed DMR (C) and DhMR (D) enrichment across comparisons. (E, F) Heatmap of DMR (E) and DhMR (F) enrichment in a region-specific manner in IPA canonical pathways altered in at least one comparison (|z-score| ≥ 1; grey, not significant). (G, H) Top IPA canonical pathways with DMR (G) and DhMR (H) enrichment in a region-specific manner altered in at least one comparison (|z-score| ≥ 1; dot, |z-score| < 1).

### Integrative analysis of RNA-seq, 5mC-seq, and 5hmC-seq data

We first examined the overlap among DEGs, DMR-associated genes, and DhMR-associated genes in the 3 comparisons to identify genes showing coordinated transcriptional and epigenetic alterations. Venn diagrams showed limited convergence among the 3 comparisons (Fig. 7A–C), suggesting that only a subset of developmental ID-induced transcriptional changes may be directly associated with alterations in both 5mC and 5hmC. In the IDA vs. IS comparison, the 4 genes that showed concurrent DEGs, DMRs, and DhMRs were primarily associated with neurodevelopment (*Pde7b*, *Tanc1*, *Cadm1*) and neuroimmune (*Cd180*) processes (Fig. 7D). In the TIDA vs. IDA comparison, the 6 genes showed coordinated concurrent DEGs, DMRs, and DhMRs were primarily associated with neurodevelopment and signaling processes (Fig. 7E).

**Figure 7.**
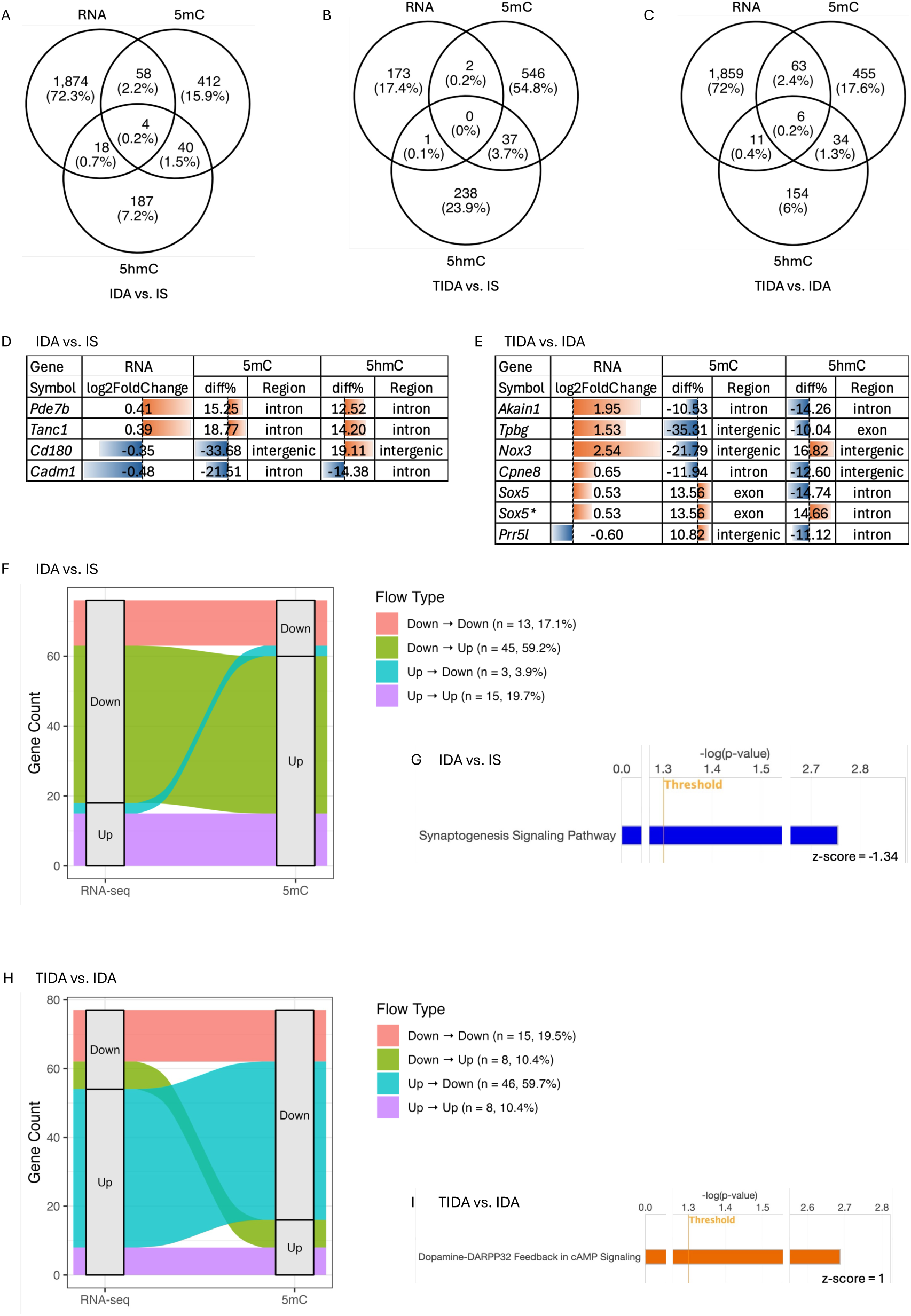
Overlap of DEGs, DMRs, and DhMRs in the 3 comparisons. (A-C) Venn diagrams showing the overlap of DEGs, DMRs, and DhMRs among the 3 comparisons. (D) The 4 genes showing concurrent differential expression, DNA methylation, and DNA hydroxymethylation in the IDA vs. IS comparison. (E) The 6 genes showing concurrent differential expression, DNA methylation, and DNA hydroxymethylation in the TIDA vs. IDA comparison. * *Sox5* contained 2 DhMRs. (F) Alluvial plot showing transitions from DEGs to 5mC enrichment at corresponding genes in the IDA vs. IS comparison. (G) IPA canonical pathway analysis of genes showing both differential expression and differential methylation in the IDA vs. IS comparison. (H) Alluvial plot showing transitions from DEGs to 5mC enrichment at corresponding genes in the TIDA vs. IDA comparison. (I) IPA canonical pathway analysis of genes showing both differential expression and differential methylation in the TIDA vs. IDA comparison.

To further investigate the relationship between gene transcription and DNA cytosine modification, we examined transitions between DEGs and corresponding DMRs and DhMRs using alluvial plots. The majority of genes (69.1–100%) showed an inverse relationship between gene transcription and DNA methylation in the 3 comparisons, with upregulated genes tending to be hypomethylated and downregulated genes tending to be hypermethylated (Fig. 7F, H; Supplemental Fig. 1). In the IDA vs. IS comparison, genes showing both differential transcription and methylation were associated with inhibited synaptogenesis signaling pathway (Fig. 7G). In the TIDA vs. IDA comparison, genes showing both differential transcription and methylation were associated with activated dopamine signaling pathway (Fig. 7I). These results suggested potential long-term dysregulation induced by developmental ID and persistent epigenetic regulation during recovery following postnatal iron treatment. The overlap between DEGs and corresponding DMRs in the TIDA vs. IS comparison, and between DEGs and corresponding DhMRs in the 3 comparisons was insufficient for pathway analysis.

A comprehensive IPA canonical pathway analysis of DEGs, DMR-associated genes, and DhMR-associated genes in the 3 comparisons showed distinct patterns of transcriptional and epigenetic regulation (Fig. 8A, B). Developmental ID induced global inhibition of biological pathways at the transcriptional level (IDA vs. IS, RNA), accompanied by increased methylation enrichment (IDA vs. IS, 5mC) in most of the corresponding pathways (Fig. 8A, B). Consistent with these patterns, transcriptome-associated pathway alterations were largely normalized in the TIDA vs. IS comparison, and methylome-associated pathway alterations showed only partial recovery (TIDA vs. IS, RNA, 5mC; Fig. 8A, B). In contrast, DhMR-associated genes showed moderate pathway enrichment in the IDA vs. IS comparison and more pronounced enrichment in the TIDA vs. IS comparison, with relatively few pathway alterations in the TIDA vs. IDA comparison (5hmC, Fig. 8A, B), suggesting a distinct pattern of 5hmC regulation. Top pathways are involved in neuronal signaling and neurodevelopment, signaling, and neuroinflammation (Fig. 8B).

**Figure 8.**
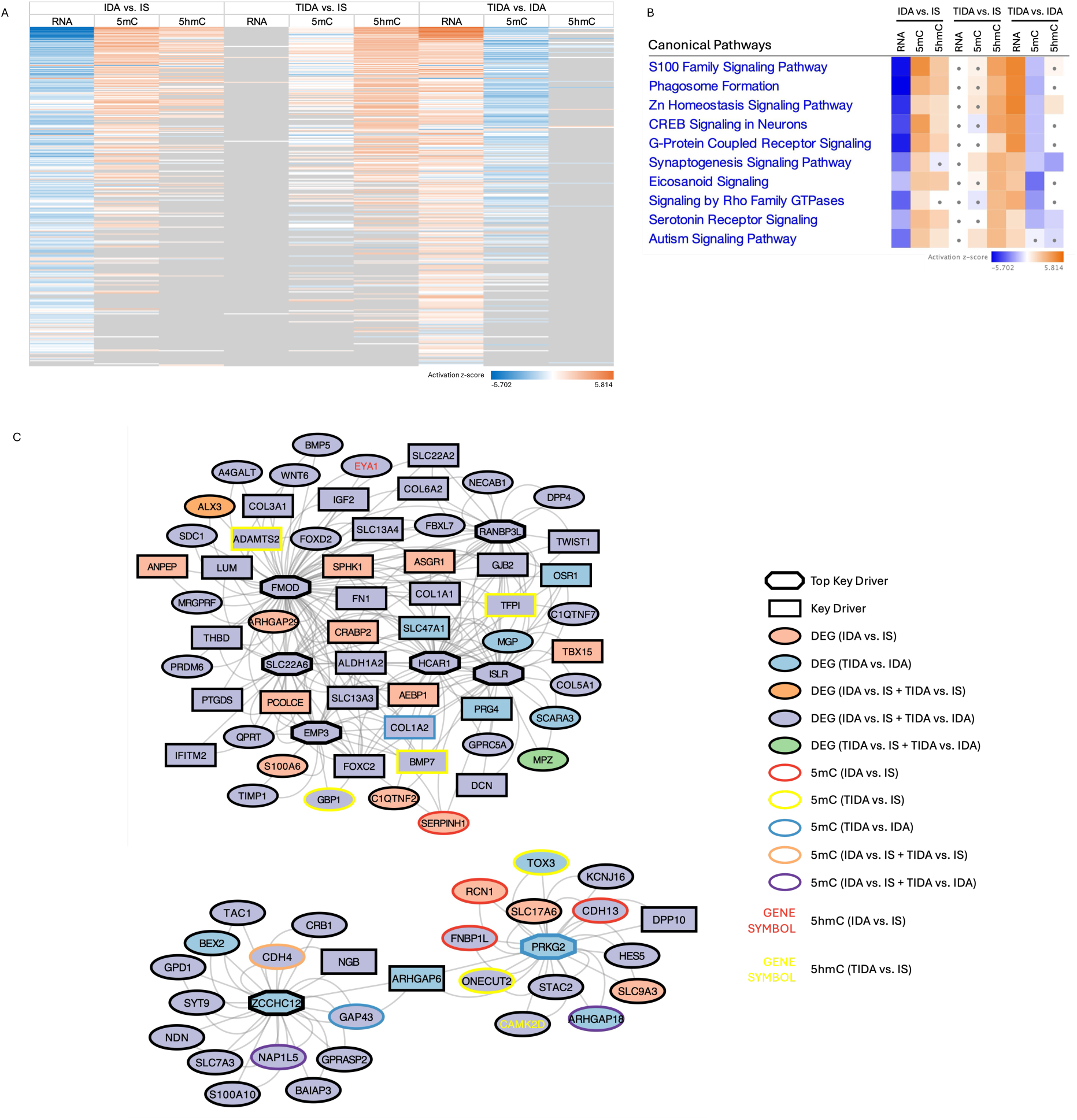
Integrative analyses of DEGs, DMRs, and DhMRs in the IDA vs. IS, TIDA vs. IS, and TIDA vs. IDA comparisons. (A) Heatmap of IPA canonical pathways altered in at least one comparison (|z-score| ≥ 1; grey, not significant). (B) Top IPA canonical pathways altered in at least one comparison (|z-score| ≥ 1; dot, |z-score| < 1). (C) MergeOmics KDA analysis identified top key drivers and the overlap with DMRs and DhMRs in the 3 comparisons.

Lastly, we performed an intergrative MergeOmics key driver analysis (KDA) to identify key drivers (KDs) in the 3 comparisons, and overlapped KDs and DEGs with DMRs and DhMRs (Fig. 8C). While most of the KDs and network genes were no longer differentially expressed following postnatal iron treatment, several remained regulated by 5mC or 5hmC (Fig. 8C, upper panel), suggesting long-term epigenetic effects of developmental ID. Notably, 3 of these genes (*Adamts2*, *Bmp7*, and *Tfpi*) were also identified as KDs, indicating persistent epigenetic regulation of central network genes despite transcriptional recovery. One top KD, *Prkg2*, showed both differential transcription and differential methylation in the TIDA vs. IDA comparison (Fig. 8C, lower panel), suggesting a potentially significant role in the molecular response to postnatal iron treatment.

## Discussion

Using multi-omics and integrative network analyses, the present study shows that developmental ID induces widespread molecular alterations in the developing hippocampus, but the molecular responses to postnatal iron treatment differ substantially across transcriptomic, 5mC, and 5hmC signatures. While transcriptional dysregulation was largely restored at P15 following iron treatment, both 5mC and particularly 5hmC exhibited complex post-treatment changes that were not explained solely by reversal of the original alterations. A large proportion of these post-treatment epigenetic alterations were new rather than residual effects of developmental ID. This pattern was particularly pronounced in 5hmC. These observations suggest that molecular recovery from developmental ID following postnatal iron treatment involves active epigenetic remodeling rather than simple reversal of prior transcriptional dysregulation. Given that iron is a cofactor for epigenetic regulators including TET and JARID protein families that regulate DNA and histone methylation (23), these findings support the broader concept that environmental insults interfering with epigenetic regulation during critical neurodevelopmental period may produce lasting molecular consequences despite immediate transcriptional recovery following removal of the insults.

The transient reversal of transcriptional changes following iron treatment suggests that gene expression in the developing hippocampus is sensitive and responsive to immediate iron availability (26). In contrast, the recovery of 5mC and 5hmC signatures were uncoupled from this transcriptomic normalization. Postnatal iron treatment reversed only a minority of the 5mC alterations. Furthermore, while developmental ID-induced 5hmC alterations exhibited a high recovery rate, this recovery does not indicate restoration of the original molecular state. Instead, a substantial proportion of post-treatment epigenetic alterations consisted of newly emerged DMRs and DhMRs. This indicates that postnatal iron treatment did not simply reverse pre-existing epigenetic dysregulation, but also resulted in extensive, *de novo* epigenetic remodeling (27) that may underlie long-term developmental consequences (24). In addition, the proportion of molecular changes directly attributable to iron treatment decreased from RNA to 5mC and further to 5hmC. This progressive decoupling suggests distinct trajectories between dynamic transcriptomic responses and more complex, persistent epigenetic regulation (37–39).

This combination of immediate transcriptional normalization and persistent epigenetic remodeling following iron treatment aligns with the paradigm of type II transcriptional memory. This paradigm has been described in the priming of mammalian immune cells (40) and plant stress responses (41); similar patterns have also been observed in the persistent metabolic changes in obesity (42). Importantly, despite limited overlap between transcriptomic and epigenetic alterations at the gene level, substantial convergence was observed at the pathway level (Fig. 7A, B), indicating network-based biological organization in which distinct molecular alterations converge on shared functional pathways (43, 44).

Developmental ID disrupts energy metabolism (45, 46), neurodevelopment (46), neurotransmitter synthesis and signaling (47), and epigenetic regulation (23, 25, 27), which may underlie long-term neurobehavioral deficits despite prompt iron treatment (48). Consistently, our pathway enrichment analysis of recovered, non-recovered, and newly emerged epigenetic alterations showed that distinct biological processes were differentially recovered or remodeled following developmental ID and postnatal iron treatment. These findings suggest a hierarchical prioritization of molecular responses. 1) Normalized modifications were primarily associated with broad biological functions, including cell migration, cell communication, and general development. In the developing rat hippocampus, the period between P7 and P15 is characterized by late-stage neurogenesis (35) and early neuronal differentiation (36). This suggests that the system prioritizes fundamental processes required for ongoing brain development over specialized functions following removal of environmental stress (i.e., ID). 2) Non-recovered modifications were notably enriched in metabolic regulation, transcriptional regulation, Wnt signaling pathway, and calcium transport. Wnt signaling is critical for adult hippocampal neurogenesis, learning, and memory (49). In addition, calcium signaling and homeostasis are essential for synaptic plasticity and long-term potentiation, which underlie learning and memory formation (50). Persistent epigenetic dysregulation of these pathways may contribute to the long-term deficits in synaptic function (51) and cognition (48) following developmental ID. 3) Newly emerged modifications were enriched in specialized pathways, including axon guidance, dopaminergic synapse, and circadian entrainment, indicating that postnatal iron treatment induces pathway-specific epigenetic remodeling beyond direct recovery. Particularly, ID in adult rats has been associated with alterations in dopamine-dependent circadian rhythm (52, 53), and ID-mediated dopamine dysregulation has been linked to Restless Legs Syndrome (54). These suggest a potential mechanism through which developmental ID may increase vulnerability to long-term disturbances in sleep and circadian regulation (55, 56).

Mechanistically, the extensive *de novo* epigenetic remodeling, particularly in 5hmC, may reflect partially recovered TET enzymatic activity following iron reintroduction (Supplemental Fig. 2). Because the TET enzymes require iron as a cofactor (57), prolonged developmental ID may directly compromise TET-mediated oxidation of 5mC to 5hmC. Consistently, we observed a global trend toward increased 5mC and decreased 5hmC levels in the IDA relative to the IS group, and a trend toward recovery in the TIDA relative to the IDA group (Supplemental Fig. 3). At the DMR and DhMR levels, the two epigenetic marks exhibited a noticeable divergence in recovery and remodeling patterns during the iron treatment period, with 5hmC showing a more dynamic pattern beyond a simple reversal of ID-induced alterations (Fig. 2D, F). This observation is consistent with the view that 5mC is a relatively stable epigenetic mark that reflects the cellular history and serves as a form of epigenetic memory (58), while 5hmC undergoes dynamic regulation during brain development and its dysregulation has been linked to cognitive and mental health disorders (59, 60). In fact, 5hmC is highly enriched in the central nervous system (61), where it plays important roles synaptic plasticity (62), memory (63), and rapid behavioral adaptation (64).

Although both DMRs and DhMRs were similarly distributed across intronic and intergenic regions, functional annotation was disproportionally associated with intronic modifications (Fig. 6). Our data showed that developmental ID primarily affects 5mC and 5hmC within non-coding genomic regions, consistent with the distribution patterns reported in other studies (65, 66). In fact, gene body 5mC has been found to be more indicative of gene expression than promoter 5mC (30, 67). Intronic 5mC regulates alternative splicing (68) and is inversely associated with gene expression (69). This may explain the concurrent suppression of pathway activity in the IDA group at the transcriptional level and increased 5mC modifications in genes associated with these pathways, driven by intronic 5mC (Figs. 6E, 7F, 8A). While non-coding region 5hmC is generally positively associated with gene expression (65), we also observed increased 5hmC modifications in genes associated with transcriptionally suppressed pathways (Fig. 8A). This finding may reflect the complex relationship between 5hmC and transcription, as previous studies have reported that 5mC and 5hmC often correlate with gene expression in the same direction (30). The coupling between DEG- and DhMR-associated pathways was also weaker than that between DEG- and DMR-associated pathways (Fig. 8A). This contrast suggests that 5hmC may reflect active epigenetic priming and reprogramming rather than directly corresponding to steady-state transcriptional activity (70). The asymmetric functional enrichment of intronic and intergenic DMRs and DhMRs likely reflects the differential genomic distribution of cis-regulatory elements (REs). Genes associated with tissue (e.g., brain)-specific functions are preferentially regulated by intronic REs, whereas housekeeping genes are more often regulated by intergenic REs (71). This may explain why primarily intronic epigenetic alterations were mapped to functionally relevant pathways.

Finally, the integrative MergeOmics analysis showed that although most of the genes altered by developmental ID were no longer differentially expressed following iron treatment, some remained epigenetically regulated and were identified as KDs. Notably, *Bmp7* is involved in neurodevelopmental signaling and neuronal differentiation (72, 73), suggesting that persistent epigenetic regulation may continue to affect developmental pathways, possibly with effects extending into adulthood (74). In addition, another top KD, *Prkg2*, responded to iron treatment at both the transcriptional and epigenetic levels. *Prkg2* is involved in cGMP-dependent signaling pathways that regulate myelination (75) and cell proliferation (76), suggesting a potential mechanism through which iron repletion restores molecular changes induced by developmental ID.

This study has several limitations. First, only male offspring were examined. We previously observed sex differences in the transcriptomic changes in the TIDA vs. IS comparison in adult (P65) rat offspring (77). This suggests that transcriptomic and epigenetic responses to developmental ID and subsequent iron treatment may also differ between sexes at P15. Second, the present study focused on P15, during active hippocampal maturation (35, 36). Although this time point provides insight into early molecular recovery and remodeling during recovery from developmental ID, it does not address whether these epigenetic alterations persist into adulthood. Our previous studies showed persistent behavioral, transcriptional, and epigenetic (chromatin accessibility, histone modification) alterations in adult male offspring (P65, TIDA vs. IS) (24–27, 77), supporting the importance of extending integrated transcriptomic, 5mC, and 5hmC analyses to later developmental stages. Future studies incorporating both sexes and additional timepoints across development will provide a more comprehensive understanding of the temporal and sex-specific dynamics of epigenetic remodeling following developmental ID and postnatal iron treatment.

## Conclusions

In this study, we investigated how prenatal ID, a nutritional perturbation that directly affects the activity of epigenetic modifiers (e.g., the TET and JARID protein families), influences transcriptional and epigenetic regulation, as well as molecular recovery following postnatal iron treatment. We found that the transcriptome showed substantial resilience following postnatal iron treatment, whereas the epigenome (5mC, 5hmC) showed only partial recovery accompanied by extensive remodeling. These findings suggest that molecular recovery following early-life environmental perturbation extends beyond normalization of gene expression and should be evaluated at the epigenetic level. Persistent molecular alterations may contribute to the long-term neurodevelopmental and behavioral deficits associated with developmental ID (23, 24). Although this study focused on developmental iron deficiency, this conceptual framework may also apply to other prenatal environmental exposures that produce enduring molecular reprogramming and neurobehavioral outcomes.

## Methods

### Animals

Timed-pregnant gestational day (G) 2 Sprague-Dawley rats (SD-400, Charles River Laboratories) were fed either iron deficient (4 mg/kg Fe, TD 80396, Envigo) or iron sufficient (200 mg/kg Fe, TD 09256, Envigo) purified diet upon arrival. The diets differed only in iron content. A random subset of litters fed the iron-deficient diet was switched to the iron-sufficient diet at postnatal day (P) 7, a time point at which rodent neurodevelopment approximates that of a human full-term infant (34). Diets were maintained until P15, when offspring were euthanized and hippocampi were collected. This generated 3 experimental groups (Fig. 1): offspring maintained on the iron-sufficient diet from G2 to P15 (IS), offspring maintained on the iron-deficient diet from G2 to P15 (IDA), and offspring maintained on the iron-deficient diet from G2 to P7 followed by iron treatment from P7 to P15 (TIDA). All litters were culled to 8 offspring (4 males, 4 females) at birth. Male offspring from at least 3 litters per experimental group were used for each analysis to minimize potential litter-specific effects (78).

Animals were housed on a 12-hour light/dark cycle with ad libitum access to food and water. Protocols were approved by the University of Minnesota Institutional Animal Care and Use Committee (Protocol # 2001-37802A).

### Hippocampal collection

Offspring were euthanized at P15 by rapid decapitation. Hippocampi were microdissected on an ice-cold metal block, flash-frozen in liquid nitrogen, and stored at -80°C until use.

### RNA extraction

Total RNA was extracted from the hippocampal tissue using the RNAqueous Total RNA Isolation Kit (Invitrogen) following the manufacturer’s instructions.

### RNA sequencing (RNA-seq) and analysis

Total RNA was treated with DNase I (Invitrogen) to remove trace amounts of genomic DNA. Library preparation and RNA-seq were performed at the University of Minnesota Genomics Center. RNA concentration was quantified using the RiboGreen RNA Assay kit (Invitrogen), and RNA quality was assessed using an Agilent 2100 BioAnalyzer (Agilent Technologies). Indexed libraries were prepared for each sample using the Illumina Stranded Total RNA Prep with Ribo-Zero Plus kit (Illumina). Libraries were size-selected (∼200 bp) and sequenced on an AVITI24 sequencer (Element Biosciences) using 150-bp paired-end reads at a depth of approximately 160 million reads per library.

Raw sequencing data quality was assessed using FastQC (v0.12.1). Reads were aligned to the *Rattus norvegicus* mRatBN7.2 reference genome using HISAT2 (v2.2.0) (79). Differentially expressed genes (DEGs) were identified using a quasi-likelihood test in edgeR (80) with thresholds of |log_2_(Fold Change)| > 0.32 (∼ 25% change) and *p* < 0.05 (n = 4/group).

### Quantitative PCR (qPCR)

RNA-to-cDNA synthesis was performed using the High-Capacity RNA-to-cDNA Kit (Applied Biosystems). qPCR was performed using the Luna Universal qPCR Master Mix (New England Biolabs) and TaqMan Gene Expression Assays (*Rps18*, Rn01428913_gH; *Tfrc*, Rn01474701_m1; *Adm*, Rn01507680_g1; *Syt8*, Rn00584120_m1; *Tmem45b*, Rn01524801_m1; Thermo Fisher Scientific) on a QuantStudio 3 Real-Time PCR System (Thermo Fisher Scientific). *Rps18* was used as the endogenous control. Samples were run in duplicate. Relative gene expression was calculated as fold change relative to the IS group using the 2^−ΔΔCt^ method. Statistical analyses were performed in Prism 11 (GraphPad Software) using one-way ANOVA followed by Holm-Šídák’s multiple comparisons test. Sample sizes ranged from n = 3–6 per group.

### DNA extraction and sequencing

Genomic DNA was extracted from hippocampal tissue using the MagAttract HMW DNA Kit (QIAGEN) following the manufacturer’s instructions. Libraries were prepared using the Ligation Sequencing Kit V14 (Oxford Nanopore Technologies, ONT). Sequencing was performed on a P2 Solo instrument (ONT) using one PromethION R10.4.1 flow cell per sample. Libraries were loaded onto the flow cell and sequenced for 24 h. Following each sequencing run, the library was recovered, the flow cell was washed using the Flow Cell Wash Kit (ONT), and the library was reloaded for the next run. Each library was sequenced over 3 consecutive runs. Data were acquired using MinKNOW software (v23.07.12, ONT) with a sampling rate of 5 kHz.

Raw data from the 3 sequencing runs were basecalled together using Dorado (v0.4.3, https://github.com/nanoporetech/dorado) with model dna_r10.4.1_e8.2_400bps_fast@v4.2.0. Reads were subsequently re-basecalled in super-accuracy mode for simultaneous 5mC and 5hmC detection using the option --modified-bases 5mC_5hmC. Read quality was assessed using Nanoq (https://github.com/esteinig/nanoq).

### ONT sequencing analysis

Global DNA methylation and hydroxymethylation (5mC and 5hmC) at cytosine-guanine dinucleotide (CpG) sites were determined using modified base information stored in the initial basecalling output files (unmapped modBAMs). Unmapped modBAMs were concatenated and aligned to the rat reference genome (mRatBN7.2) using Minimap2 (81). The resulting mapped modBAMs were converted to bedMethyl format using Modkit (v0.2.2, https://github.com/nanoporetech/modkit). Differential analyses of 5mC and 5hmC (DMR and DhMR) were performed using MethylLasso (82) with minimum coverage of 5 reads per CpG site and 4 CpG sites per region. DMRs were identified using a minimum methylation difference threshold of 10% and a false discovery rate (FDR) threshold of *q* < 0.05 (n = 3/group). DhMRs were identified using a minimum methylation difference threshold of 10% and a significance threshold of *p* < 0.05 (n = 3/group) (83, 84).

DMRs and DhMRs were annotated by genomic feature and nearest gene using the mRatBN7.2.111 GTF file. Promoters were defined as 2 kb upstream to 500 bp downstream of transcription start sites, and regions were classified as promoter, exon, intron, or intergenic using the priority order promoter > exon > intron > intergenic.

### Gene Oncology (GO) and Kyoto Encyclopedia of Genes and Genomes (KEGG) analyses

GO and KEGG analyses were performed using ShinyGO (v0.85.1) (85). The FDR threshold was set at 0.05, redundancy was removed, and the top 10 enriched terms were reported.

### Ingenuity Pathway Analysis (IPA)

DEGs, DMR- and DhMR-associated genes were analyzed using QIAGEN IPA software (QIAGEN) to identify significantly altered canonical pathways and Diseases & Bio Functions. Statistical significance (*p* < 0.05; -log(*p*) > 1.3) was determined using Fisher’s exact test, and pathway activation states were predicted using |z-score| ≥ 1.

### Integrative network analysis

Integrative network analysis was performed using the Mergeomics pipeline (86). DEGs from all comparisons were integrated as separate modules into a preconstructed brain-specific gene regulatory network, and key driver analysis (KDA) was used to identify candidate key regulatory genes (key drivers, KDs) based on significant local network enrichment. DMR- and DhMR-associated genes were subsequently overlaid onto the resulting KDA networks to identify epigenetically regulated network genes and KDs.

### Nuclear protein isolation

Nuclear proteins were isolated from the hippocampus using the EpiQuik Nuclear Extraction Kit II (EpigenTek Group Inc.) following the manufacturer’s instructions. Nuclear protein concentration was quantified using a Bradford protein assay.

### TET1 activity

TET1 enzymatic activity was assessed using nuclear protein extract with the Epigenase 5mC-Hydroxylase TET Activity/Inhibition Assay kit (EpigenTek Group Inc.). Outliers were identified and removed using the 1.5 x interquartile range method. Statistical analyses were performed in Prism 11 (GraphPad Software) using one-way ANOVA followed by Holm-Šídák’s multiple comparisons test. Sample sizes ranged from n = 4–7 per group.

## Abbreviations

CpG: Cytosine-guanine dinucleotide
DEG: Differentially expressed gene
DhMR: Differentially hydroxymethylated region
DMR: Differentially methylated region
FDR: False discovery rate
G: Gestational day
GO: Gene ontology
ID: Iron deficiency
IDA: Iron-deficient anemic
IPA: Ingenuity Pathway Analysis
IS: Iron-sufficient
JARID: Jumonji and AT-Rich Interaction Domain-containing
KD: Key driver
KDA: Key driver analysis
KEGG: Kyoto Encyclopedia of Genes and Genomes
NS: Not significant
ONT: Oxford Nanopore Technologies
P: Postnatal day
qPCR: Quantitative PCR
TET: Ten-Eleven Translocation
TIDA: Treated IDA

## Declarations

## Availability of data and materials

The datasets generated and analyzed during the current study will be uploaded to the Gene Expression Omnibus repository.

## Authors’ contributions

SXL and PVT conceptualized and designed the work. SXL, ZLM, CW, CK, and CF performed the experiments and acquired the data. SXL, CK, and CF analyzed and interpreted the data. SXL drafted the manuscript. SXL, MKG, and PVT substantively revised the manuscript. All authors read and approved the final manuscript.

## Acknowledgements

We thank Dr. Natalie Calixto Mancipe for assistance with the initial RNA-seq analysis.

