## Supplementary figures and images for "Hierarchical Transcriptomic and Epigenetic Recovery and Remodeling in the Developing Hippocampus Following Early-life Environmental Insults: An Iron Deficiency Rat Model"

### Supplemental Figure 1

#### Slide 1
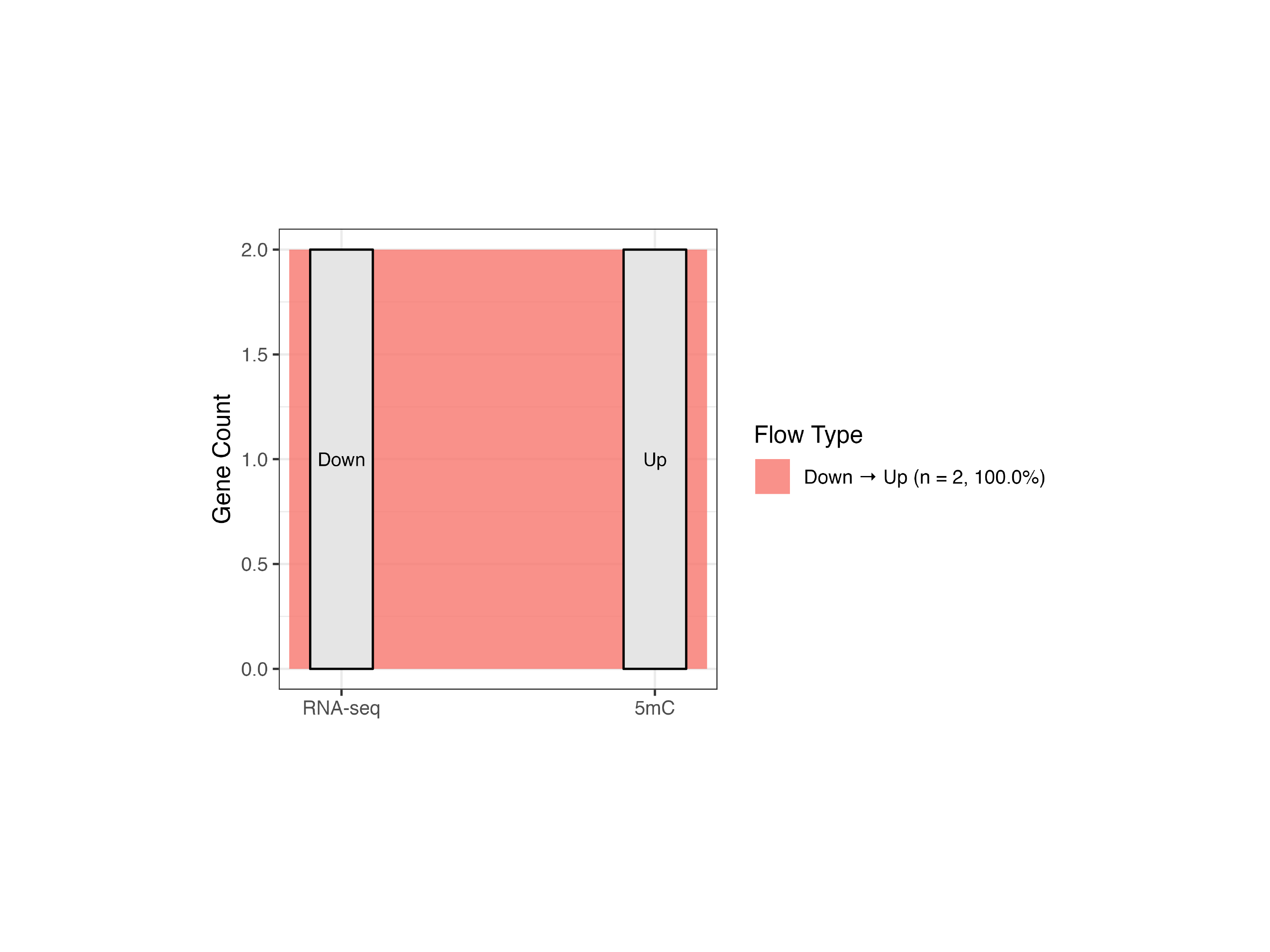

### Supplemental Figure 2

#### Slide 1
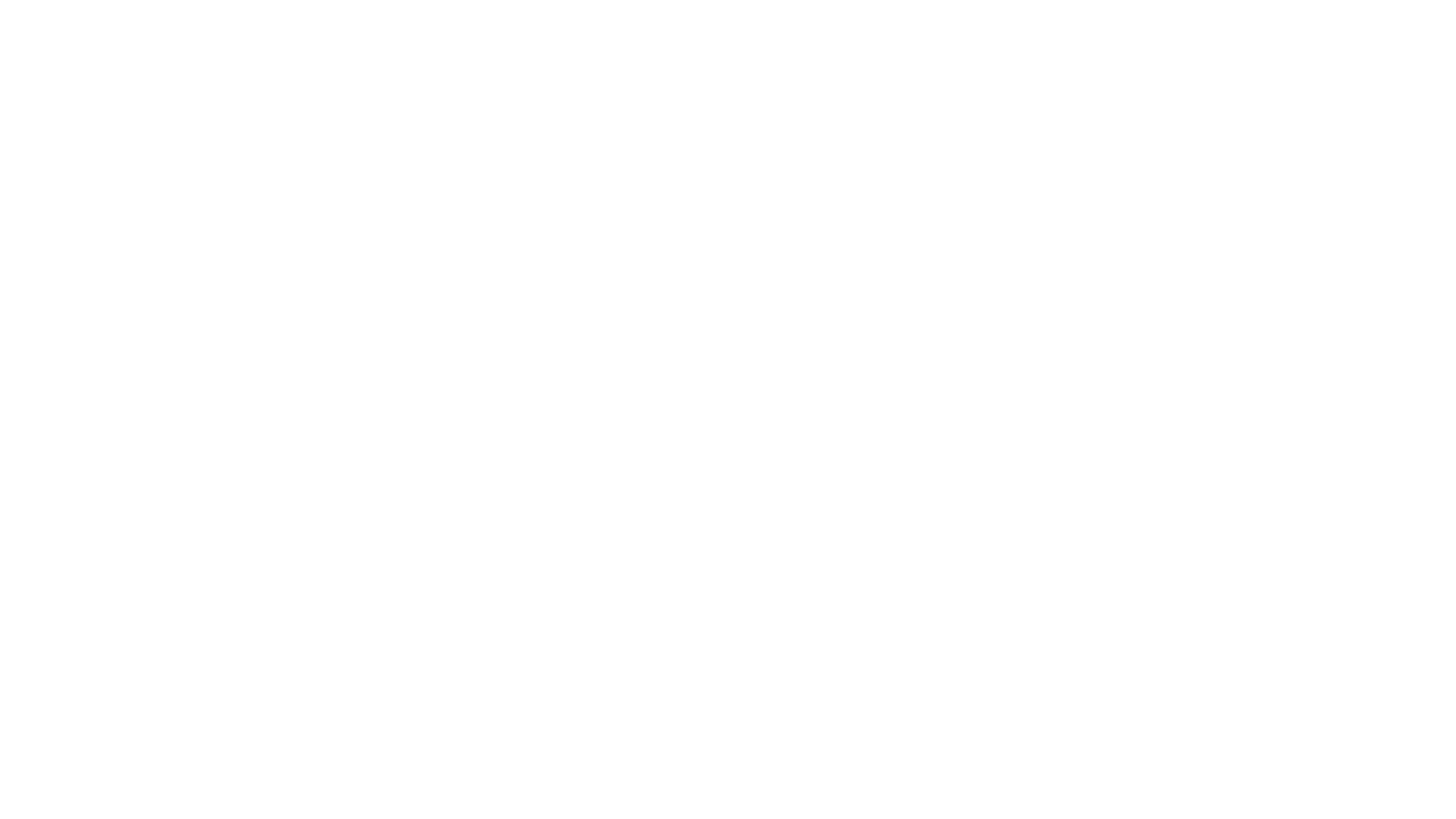

### Supplemental Figure 3

#### Slide 1
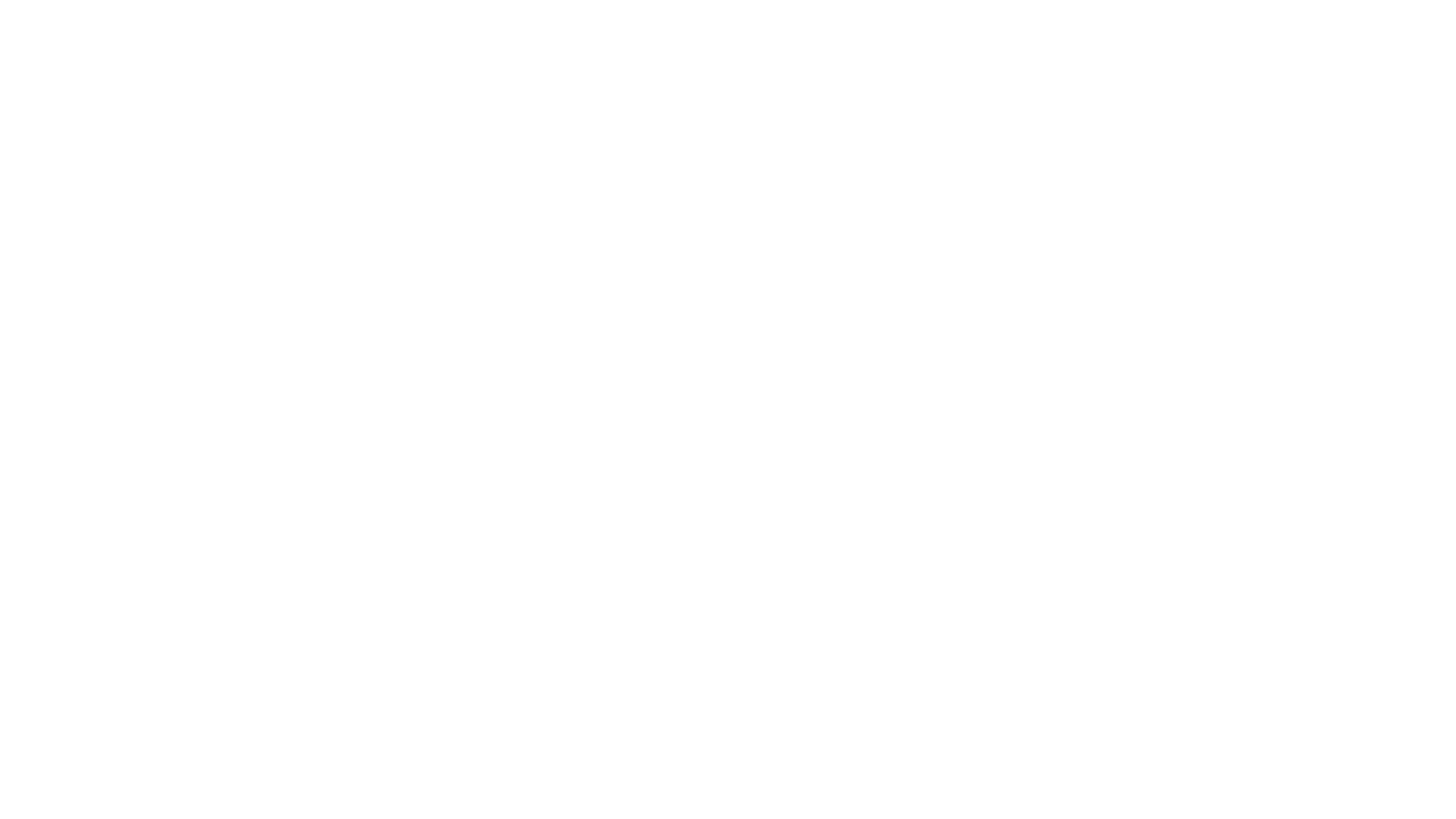
